# Statin-dye conjugates for selective targeting of *KRAS* mutant cancer cells

**DOI:** 10.1101/2025.06.03.657329

**Authors:** Hye-ran Moon, Zhenying Cai, Bo Kyung Cho, Hyeyoun Chang, Seung Taek Hong, Jean J. Zhao, Ick Chan Kwon, Thomas M. Roberts, Bumsoo Han, Ju Hee Ryu

## Abstract

Over 90% of pancreatic ductal adenocarcinoma (PDAC) patients involve *KRAS* mutations (*KRAS*^MUT^), for which current treatment options are limited. Statins, commonly used to lower cholesterol, have demonstrated certain selective toxicity towards *KRAS*-transformed cells, prompting the question of whether statins could achieve selective uptake specifically in *KRAS*^MUT^ cells. To investigate this, we synthesized statin-dye conjugates by attaching a fluorescent dye (Cy5.5) to two statins: simvastatin and pravastatin, aiming to assess whether selective uptake indeed occurs. Our findings revealed that these conjugates exhibited markedly enhanced uptake in *KRAS*^MUT^ cells compared to *KRAS* wild-type (*KRAS*^WT^) cells. Given the magnitude of the selective uptake, we realized that the uptake of these conjugates itself is of considerable intrinsic interests. We evaluated the uptake of these conjugates in both *KRAS*^MUT^ and *KRAS*^WT^ cells and examined their potential to selectively target *KRAS*^MUT^ pancreatic cancer cells (PCCs) using an engineered PDAC tumor model co-cultured with PCCs and cancer-associated fibroblasts (CAFs). Our findings indicate that *KRAS*^MUT^ cancer cells exhibited higher uptake of statin-Cy5.5 conjugates *via* enhanced macropinocytosis compared to *KRAS*^WT^ cancer cells and CAFs. We also found enhanced uptake of the statin-Cy5.5 conjugate in MCF10A cells with *PTEN* deficiency, a condition known to elevate macropinocytosis, compared to control MCF10A cells with wild-type *PTEN*. Notably, in the PCC and CAF co-culture model, the pravastatin-Cy5.5 conjugate selectively killed *KRAS*^MUT^ PCCs without affecting the *KRAS*^WT^ CAFs. These findings highlight the potential of stain-drug conjugates as targeted delivery vehicles for *KRAS*^MUT^ cancer therapy.

## INTRODUCTION

Pancreatic ductal adenocarcinoma (PDAC) is the third leading cause of cancer-related death in the United States, with a five-year survival rate of only 12%.^1^ PDAC’s aggressive nature and resistance to conventional treatments, such as chemotherapy, radiotherapy, targeted therapy, and immunotherapy, underscore the urgent need for new therapeutic approaches. The tumor microenvironment (TME) of PDAC is particularly complex, consisting of pancreatic cancer cells (PCCs), cancer-associated fibroblasts (CAFs), and various immune cells, further complicating treatment.^2^ Consequently, current treatment options are often inadequate, and novel strategies are needed to improve patient outcomes.

A key feature of PDAC is the high prevalence of *KRAS* mutations, present in over 90% of PDAC patients.^3^ These mutations play a crucial role in driving tumorigenesis by maintaining *KRAS* in a continuously active state, promoting uncontrolled cell growth and division. This persistent signaling contributes to the cancer’s aggressive behavior and poor prognosis, making *KRAS* a significant target for cancer therapy. Recent breakthroughs have shown promise; for example, the *KRAS*^G12C^ inhibitor sotorasib (AMG 510) demonstrated promising results in phase I clinical trial for non-small cell lung cancer, with partial response or stable disease observed in 88.1% of patients with the *KRAS*^G12C^ mutations.^4, 5^ Additionally, a recent study suggests that *KRAS*^G12D^ inhibitors may offer potential benefits in treating pancreatic cancers.^6^ However, the heterogeneity of *KRAS* mutations, including subtypes including G12C, G13D, G12D, G12V, and G12R presents a challenge for developing treatments that are effective across all variants.^7^ Therefore, there is an urgent need to develop therapies that can effectively target the diverse range of *KRAS* mutation subtypes.

Interestingly, statins, widely used to lower cholesterol, have shown efficacy in killing *KRAS* mutant (*KRAS*^MUT^) cells *in vitro* and in tumor models.^5, 8–10^ Statins inhibit the mevalonate pathway, necessary for *KRAS* prenylation, a post-translational modification required for its localization to the cell membrane where it exerts oncogenic effects.^9, 11–14^ By inhibiting prenylation, statins disrupt *KRAS* function, impeding its role in promoting tumor growth.^15^ This raises the question of whether statins might also achieve selective uptake specifically in *KRAS*^MUT^ cells.

To investigate this, we synthesized statin-dye conjugates by attaching a fluorescent dye (Cy5.5) to two statins, simvastatin and pravastatin, aiming to assess whether selective uptake occurs. To our knowledge, no previous studies have directly examined selective uptake of statins with fluorescent labeling in *KRAS*^MUT^ cells, and we sought to investigate the uptake mechanism of these conjugates. Our findings revealed that these conjugates exhibited markedly enhanced uptake in *KRAS*^MUT^ cells compared to *KRAS* wild-type (*KRAS*^WT^) cells. Given the magnitude of the selective uptake, we realized that the uptake of these conjugates itself is of considerable intrinsic interests.

Furthermore, we investigated their ability to selectively target *KRAS*^MUT^ PCCs using an engineered PDAC tumor model with co-cultures of PCCs and CAFs. Our results showed that these statin-dye conjugates were selectively taken up by *KRAS*^MUT^ cells as compared to isogenic *KRAS*^WT^ cells and CAFs. We also found that statin-Cy5.5 conjugates were selectively taken up in *PTEN*^KO^ cells *via* macropinocytosis. Moreover, the pravastatin-Cy5.5 conjugate showed modest selective killing toward PCCs in a 3D co-culture model. This suggests that statins have the potential for selective uptake in *KRAS*^MUT^ cells, a characteristic that could drive the development of stain-drug conjugates as targeted therapies for *KRAS*^MUT^ cancers. These findings highlight the potential of stain-drug conjugates as targeted delivery vehicles for *KRAS*^MUT^ cancer therapy.

## RESULTS

### Synthesis of statin-dye conjugates

We synthesized statin-dye conjugates by attaching the fluorescent dye Cy5.5 to simvastatin and pravastatin. Simvastatin was conjugated to Cy5.5 using 1-ethyl-3-(3-dimethylaminopropyl) carbodiimide (EDC) and 4-dimethylaminopyridine (DMAP), resulting in the formation of simvastatin-Cy5.5. Pravastatin was similarly conjugated with Cy5.5 using EDC and N-hydroxysuccinimide (NHS), producing pravastatin-Cy5.5 (Figure 1a). The synthesized conjugates, simvastatin-Cy5.5 and pravastatin-Cy5.5, were analyzed using high-performance liquid chromatography (HPLC) and mass spectrometry (Figure 1b,c). Mass spectrometry confirmed the expected molecular weights of the conjugates, validating successful synthesis. HPLC analysis further confirmed the purity of both conjugates, achieving over 95% purity.

**Figure 1.**
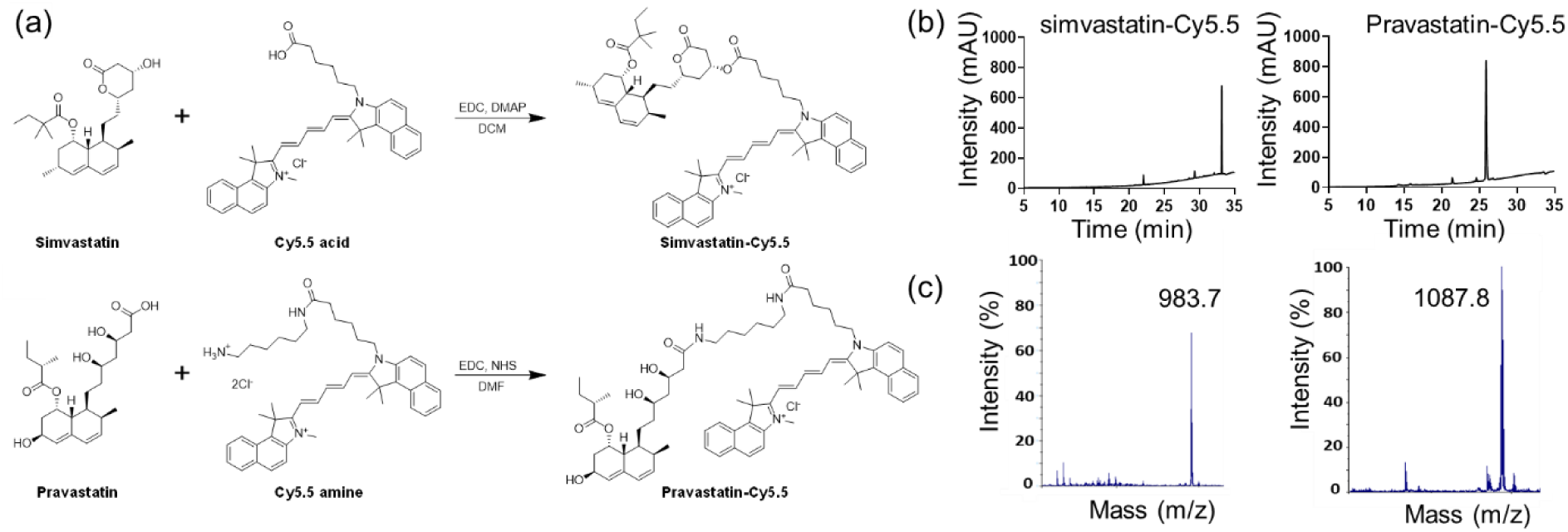
Synthesis and characterization of the statin-dye conjugates. (a) Schematic diagram illustrating the synthesis of the simvastatin-Cy5.5 and pravastatin-Cy5.5 conjugates. EDC: 1-ethyl-3-(3-dimethylaminopropyl) carbodiimide, DMAP: 4-dimethylaminopyridine, DCM: dichloromethane, NHS: N-hydroxysuccinimide, DMF: dimethylformamide. (b) High-performance liquid chromatography (HPLC) chromatograms and (c) mass spectrometry analysis of simvastatin-Cy5.5 and pravastatin-Cy5.5.

### Selective uptake of statin-dye conjugates in *KRAS*^MUT^ cancer cells

To investigate the preferential uptake of statin-dye conjugates in *KRAS*^MUT^ cancer cells, we evaluated the cellular uptake of simvastatin-Cy5.5, pravastatin-Cy5.5, and Cy5.5 across several cell lines. We first compared the uptake of these statin-dye conjugates in two distinct pancreatic cancer cell types: Panc1, featuring a *KRAS* G12D mutation (*KRAS*^G12D^), and BxPC3 with *KRAS*^WT^ using confocal microscopy. Notably, the uptake of simvastatin-Cy5.5 was significantly higher in Panc1 cells than in BxPC3 cells (Figure 2a). Similarly, the uptake of pravastatin-Cy5.5 was also significantly higher in Panc1 cells compared to in BxPC3 cells, although the level of uptake was lower than that of simvastatin-Cy5.5 in Panc1 cells. Free Cy5.5 also showed selective uptake by *KRAS*^MUT^ cancer cells *in vitro*. To further assess the concentration dependency of statin-dye conjugate uptake *via* macropinocytosis, we examined the uptake of simvastatin-Cy5.5 and Cy5.5 alone in KRAS^MUT^ Panc1 cells at two different concentrations: 50 nM and a lower concentration of 17 nM (1/3 of 50 nM). Our results demonstrate that simvastatin-Cy5.5 exhibits concentration-dependent cellular uptake in Panc1 cells, with a higher intracellular accumulation at 50 nM compared to 17 nM. In contrast, Cy5.5 alone showed minimal uptake at both concentrations, suggesting that the statin moiety plays a crucial role in facilitating uptake in KRAS^MUT^ cells (Figure S1). Flow cytometric data quantitatively assessing the uptake of the statin-dye conjugates indicated significantly enhanced uptake of both simvastatin-Cy5.5 and pravastatin-Cy5.5 compared to Cy5.5 alone (Figure S2a).

**Figure 2.**
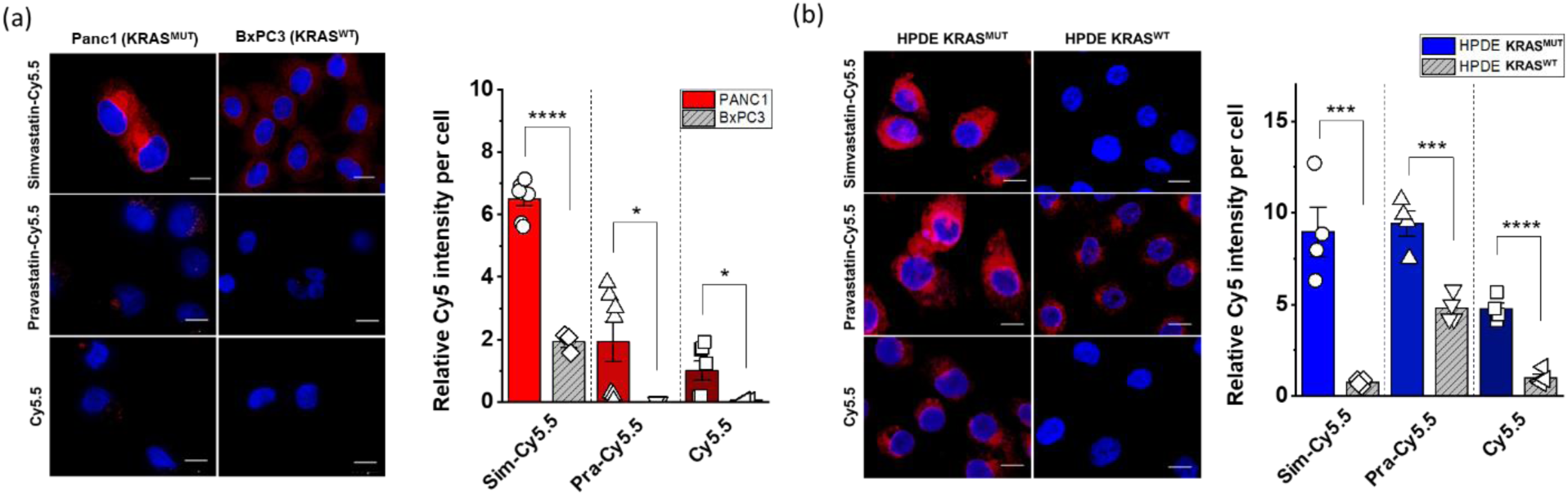
Cellular uptake of statin-dye conjugates by *KRAS*^MUT^ and *KRAS*^WT^ cells. Simvastatin-Cy5.5, pravastatin-Cy5.5, and free Cy5.5 uptake was measured by fluorescence (red) in (a) Panc1 (*KRAS*^MUT^) and BxPC3 (*KRAS*^WT^) and (b) HPDE *KRAS*^MUT^ (HPDE i*KRAS* with doxycycline) and HPDE *KRAS*^WT^ (HPDE i*KRAS* without doxycycline). The scale bars indicate 10 μm. The cell nuclei were stained with DAPI (blue). Quantitative measurements of the cellular uptake of the statin-dye conjugates are presented relative fluorescence intensity per cell count. Bars indicate Mean ± S.E. (n ≥ 3). Statistically significant differences are represented as * for p < 0.05, *** for p < 0.001, and **** for p < 0.0001.

To further evaluate the potential preferential uptake of statin-dye conjugates in *KRAS*^MUT^ cells, we assessed their uptake in genetically modified human pancreatic ductal epithelial (HPDE) cells engineered to inducibly express activated *KRAS*^G12D^ following doxycycline treatment (HPDE *iKRAS*).^16^ Notably, we observed significantly higher fluorescence intensity from the statin-dye conjugates in HPDE *iKRAS* cells treated with doxycycline induction (HPDE *KRAS*^MUT^) than in control cells without doxycycline induction (HPDE *KRAS*^WT^) (Figure 2b). Quantitative assessment of the relative Cy5.5 intensity per cell revealed a significantly augmented accumulation of all compounds, simvastatin-Cy5.5, pravastatin-Cy5.5 and Cy5.5 in *KRAS*^MUT^ cells as opposed to *KRAS*^WT^ cells. Additionally, to determine whether the selective uptake of statin-Cy5.5 conjugates observed in *KRAS*^G12D^ cells extends to other KRAS mutation subtypes, we tested two isogenic colorectal cell line pairs, HCT116 and DLD1, which carry the *KRAS* G13D mutation alongside their *KRAS*^WT^ counterparts (Table S1). Consistent with the results from *KRAS*^G12D^ cells, we observed significantly higher uptake of the statin-Cy5.5 conjugates in *KRAS*^WT/G13D^ cells compared to their *KRAS*^WT^ controls (Figure S2b,c). These findings suggest that statin-Cy5.5 conjugates undergo specific uptake in *KRAS*^MUT^ cells, indicating a potentially distinct transport mechanism for these statin conjugates in *KRAS*^MUT^ cells.

### Macropinocytosis as a key mechanism in statin-dye conjugate uptake

Given that macropinocytosis is known to be activated in *KRAS*^MUT^ cells to facilitate nutrient uptake,^17^ we next tested whether this process might contribute to the enhanced uptake of stain-Cy5.5 conjugates. Macropinocytosis is a form of endocytosis that allows cells to engulf extracellular fluid and molecules, which is often upregulated in cancer cells with certain mutations, including *KRAS* mutations. To test this hypothesis, we employed the pharmacological inhibitor 5-(N-ethyl-N-isopropyl)amiloride (EIPA), which specifically targets macropinocytosis. Before assessing the effects of EIPA on statin-dye conjugates uptake, we first validated its efficacy in inhibiting macropinocytosis using fluorescein isothiocyanate-labeled BSA (FITC-BSA), a widely recognized marker for macropinocytosis (Figure S3). *KRAS*^MUT^ Panc1 cells were pre-treated with 50 μM EIPA for 1.5 hours, followed by a 1-hour incubation with FITC-BSA (2 mg/mL). As expected, EIPA treatment significantly reduced FITC-BSA uptake, confirming its effectiveness as a macropinocytosis inhibitor. Additionally, we also confirmed that *KRAS*^MUT^ Panc1 cells exhibited higher macropinocytotic activity compared to *KRAS*^WT^ BxPC3 cells, further supporting the selective engagement of this pathway in *KRAS*^MUT^ cells using FITC-BSA.

Following this validation, we conducted EIPA experiments on statin-dye conjugates to determine whether their uptake was mediated by macropinocytosis. As shown in Figure 3a, the intracellular fluorescence from both simvastatin-Cy5.5 and pravastatin-Cy5.5 was significantly decreased when EIPA was added to the *KRAS*^MUT^ cells in both Panc1 and HPDE *KRAS*^MUT^-induced cells. . The selective uptake of statin-dye conjugates in *KRAS*^MUT^ cells, along with the significant reduction in intracellular fluorescence following EIPA treatment, indicates that macropinocytosis plays a significant role in their uptake in *KRAS*^MUT^ cells.

**Figure 3.**
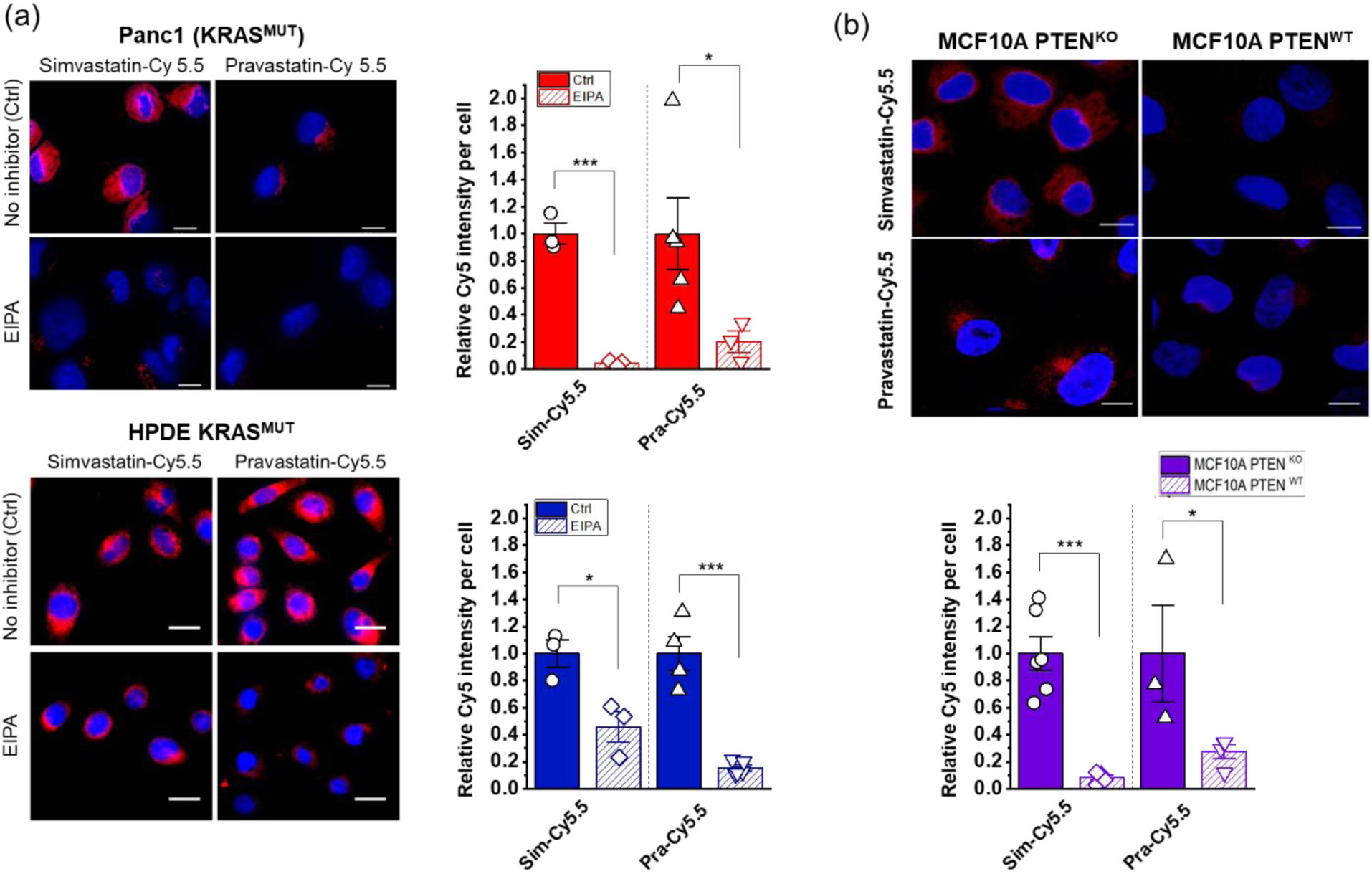
Regulation of cellular uptake of statin-Cy5.5 conjugates by inhibitors and pathways associated with macropinocytosis. (a) Treatment with 5-(N-ethyl-N-isopropyl)amiloride (EIPA) blocks the selective uptake of statin-dye conjugates in *KRAS*^MUT^ cells, specifically Panc1 and HPDE *KRAS*^MUT^. Ctrl (Control) indicates cells with no EIPA treatment. (b) MCF10A cells with *PTEN*^KO^ show enhanced uptake of stain-dye conjugates (red) compared to *PTEN*^WT^ MCF10A control cells. The cell nuclei were stained with DAPI (blue). The scale bars indicate 10 μm. Bars represent Mean ± S.E. (n ≥ 3). Statistically significant differences are represented as * for p < 0.05 and *** for p < 0.001.

We also measured the uptake of statin-dye conjugates in cells with *PTEN* deficiency, a condition known to elevate macropinocytosis due to the loss of phosphoinositide 3-kinase (PI3K) inhibition. *PTEN* is a tumor suppressor gene that negatively regulates the PI3K/AKT signaling pathway. Loss of *PTEN* function leads to increased activation of the PI3K pathway, which can promote macropinocytosis, a process often upregulated in cancer cells to meet their increased nutrient demands. By using CRISPR-Cas9 technology, we created isogenic cell lines with either sgRNA targeting *PTEN* (*PTEN*^KO^) or sgRNA targeting a non-coding sequence (*PTEN*^WT^) in MCF10A cells. As can be seen in Figure 3b, MCF10A cells with *PTEN*^KO^ exhibited stronger fluorescence compared to the *PTEN*^WT^ isogenic control, indicating enhanced uptake of simvastatin-Cy5.5 and pravastatin-Cy5.5. Upon treatment with either EIPA or a pan-PI3K inhibitor (BKM120), the fluorescence signal of simvastatin-Cy5.5 in MCF10A *PTEN*^KO^ cells decreased significantly (Figure S4). These results suggest that *PTEN* loss can increase the cellular uptake of statin-Cy5.5 conjugates in a PI3K pathway-dependent manner, further highlighting the role of macropinocytosis in this process.

### Selective cytotoxicity of statin-dye conjugates in *KRAS*^MUT^ cells *in vitro*

Statins have been reported to kill *KRAS*^MUT^ cancer cells more effectively than *KRAS*^WT^ cells.^8, 9^ To investigate the selective cytotoxic effects of statin-Cy5.5 conjugates on *KRAS*^MUT^ cancer cells, we performed cytotoxicity assays using HPDE *iKRAS* cell monolayers. Pravastatin-Cy5.5 displayed some selective effectiveness against HPDE *KRAS*^MUT^ cells compared to HPDE *KRAS*^WT^ cells at concentrations of 0.05 µM and 0.1 µM (Figure 4a). However, neither simvastatin-Cy5.5 nor Cy5.5 alone demonstrated selective killing effects on HPDE *KRAS*^MUT^ cells compared to HPDE *KRAS*^WT^ cells. Additionally, in MCF10A *PTEN*^KO^ cells that exhibited selective uptake of statin-Cy5.5 conjugates, no significant differences in cytotoxic effects were observed compared to their isogenic *PTEN*^WT^ counterparts (Figure 4b). This narrow range of efficacy urges a more comprehensive investigation into the underlying mechanisms.

**Figure 4.**
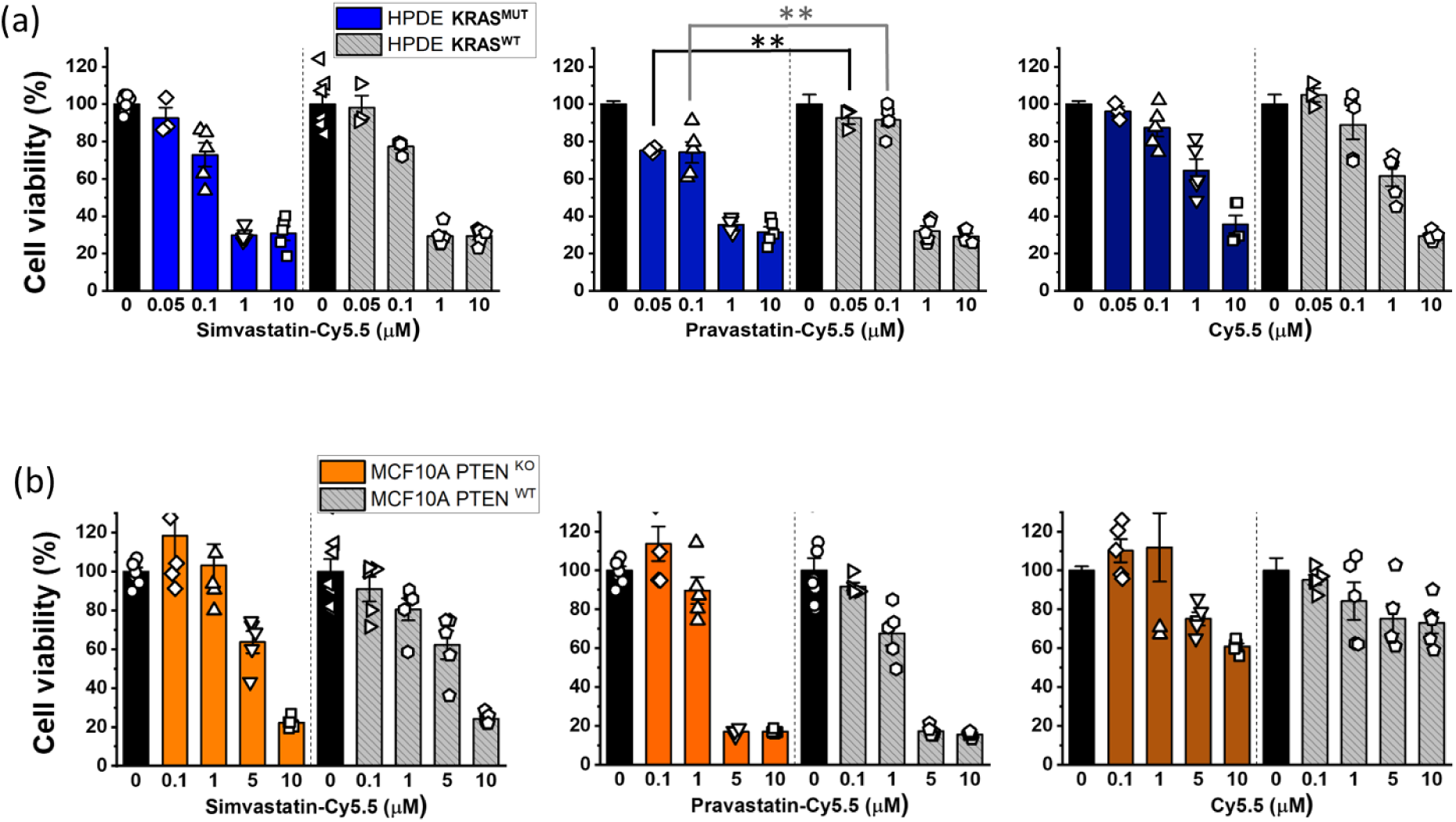
The cytotoxic effects of statin-dye conjugates. Cell viability (%) was measured after treating the 2D cell monolayers with simvastatin-Cy5.5, pravastatin-Cy5.5, and free Cy5.5 in (a) HPDE *KRAS*^MUT^ and *KRAS*^WT^ cells and (b) MCF10A with *PTEN*^KO^ and MCF10A with *PTEN*^WT^. Bars represent Mean ± S.E. (n ≥ 3). Statistically significant differences with p < 0.01 are indicated by **.

### *KRAS*^MUT^ cancer cells selectively uptake statin-Cy5.5 conjugates in the engineered TME

To further evaluate the transport and accumulation of statin-Cy5.5 conjugates in targeting *KRAS*^MUT^ cancer cells, we expanded our assessment using a microfluidic TME-on-a-chip model (T-MOC). This biomimetic tumor model recapitulates the stroma tissue in PDAC, where PCCs and CAFs are embedded in a 3D matrix. The fluid flow passes through an endothelium-mimicking membrane interfaced with a capillary channel (Figure 5a), simulating the hydrodynamic conditions of drug transport. The T-MOC model is designed to regulate the transport of fluid, nutrients, and drugs by adjusting hydrostatic pressure variations across channels, mimicking the dynamic transport conditions within the TME.^18–21^ Additionally, this system reconstitutes pharmacokinetic processes including extravasation from a capillary vessel, interstitial diffusion and convection, cellular uptake, and lymphatic drainage.

**Figure 5.**
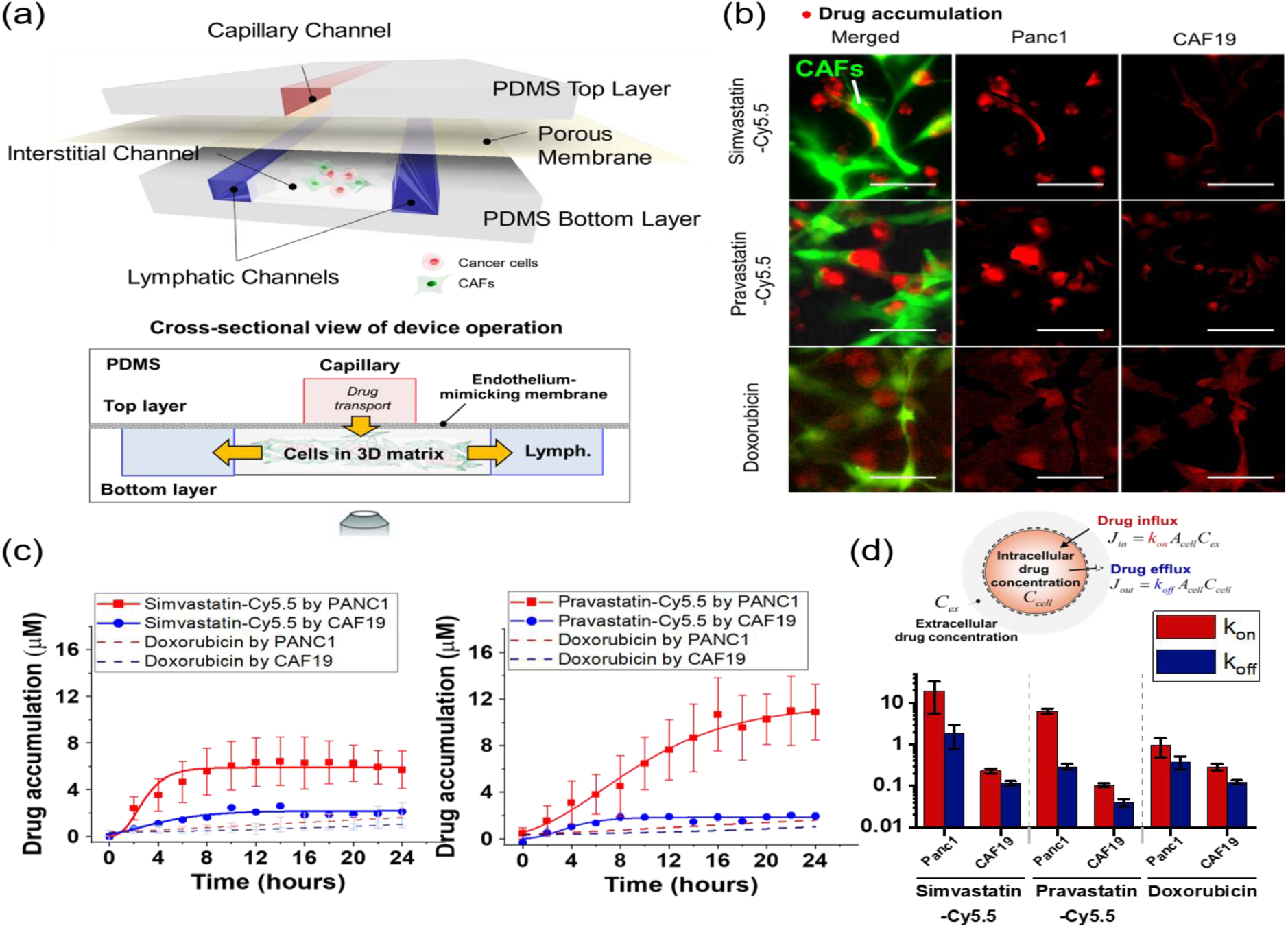
Selective uptake of statin-Cy5.5 conjugates into *KRAS*^MUT^ Panc1 cells over *KRAS*^WT^ cancer-associated fibroblasts (CAFs) in an *in vitro* cancer-stroma tumor environment-on-a-chip model (T-MOC). (a) Schematic configuration of the cancer-stroma T-MOC model operation, illustrating the setup and flow dynamics within the model. (b) Micrograph showing drug accumulation of simvastatin-Cy5.5, pravastatin-Cy5.5, and doxorubicin (used as a control). In the image, red indicates accumulated drugs, and green indicates transfected CAFs. The scale bars represent 100 μm. (c) Quantified drug accumulation measured in cell type-specific areas for simvastatin-Cy5.5, pravastatin-Cy5.5, and doxorubicin. Bars indicate Mean ± S.E. (n ≥ 3). (d) Drug accumulation model based on mass conservation principles, illustrating the quantified rate constants of drug influx and efflux within the T-MOC system.

Using the T-MOC model, we perfused either simvastatin-Cy5.5 or pravastatin-Cy5.5 along the capillary channel and monitored transient drug accumulation with time-lapse microscopy. All T-MOC experiments were conducted using living cells without fixation, allowing us to assess cellular uptake under physiologically relevant conditions. The PDAC tumor model with T-MOC included non-transfected cancer cells (Panc1) and green fluorescent protein (GFP)-transfected CAFs (CAF19), allowing for clear differentiation of drug accumulation in each cell type. Results showed that the fluorescence intensity of simvastatin-Cy5.5 and pravastatin-Cy5.5 was significantly higher in the Panc1 cell area compared to the CAF19 area. In contrast, doxorubicin, used as a control drug, showed no significant difference in fluorescence intensity between Panc1 and CAF19 cells (Figure 5b). This comparison highlights the preferential uptake of statin-Cy5.5 conjugates in *KRAS*^MUT^ cancer cells over CAFs, unlike doxorubicin.

Drug accumulation was measured by calibrating fluorescence intensity to 2 μM concentrations of simvastatin-Cy5.5, pravastatin-Cy5.5, and doxorubicin within the capillary channel. Quantitative results showed that *KRAS*^MUT^ Panc1 cells exhibited significantly higher uptake of both simvastatin-Cy5.5 and pravastatin-Cy5.5 compared to CAF19 cells (Figure 5c). Specifically, simvastatin-Cy5.5 accumulated approximately 2.5 times more, and pravastatin-Cy5.5 around 5 times more, in Panc1 cells compared to CAF19 cells during a 24 h perfusion. In contrast, doxorubicin showed no significant difference in accumulation between the two cell types (dash lines in Figure 5c), suggesting that statin-Cy5.5 conjugates selectively target cancer cells with minimal impact on surrounding stromal cells in PDAC tissue.

To further investigate drug transport mechanisms, we applied a simple drug accumulation model based on mass conservation (Figure 5d)^21^. In this model, the parameter k_on_ represented the cellular capability for drug uptake affinity at the cell surface, while k_off_ represented drug efflux as the drug dissociation affinity. Assuming that k_on_ and k_off_ are constant, quantitative analysis showed that k_on_ values for Panc1 cells were significantly higher than those for CAF19 cells for both statin-Cy5.5 conjugates. Although Panc1 cells also exhibited a higher k_on_ for doxorubicin than CAF19, this difference was less pronounced compared to statin-Cy5.5 conjugates. The high k_on_ to k_off_ ratio for statin-dye conjugates in *KRAS*^MUT^ Panc1 cells suggests that selective uptake may be linked to cancer cell-specific endocytic activity, particularly macropinocytosis, which is known for facilitating large-scale uptake.

### Tumor-selective cytotoxicity of statin-dye conjugates using the engineered TME

Building on previous findings that demonstrated the *KRAS*^MUT^ targeting potential of statin-dye conjugates, we hypothesized that these conjugates could selectively kill *KRAS*^MUT^ tumor cells over *KRAS*^WT^ stroma cells in tumor tissue. To maximize screening effectiveness, we applied the drug solution in a stationary manner, eliminating flow dynamics. Devices were pre-cultured for 48 h, followed by a 24-h drug treatment. After treatment, the model was perfused with drug-free media and cultured for an additional 48 h to capture any latent drug effects. Cell viability was assessed by measuring the stained nuclear area of the non-transfected PCCs (Panc1) and GFP-transfected CAFs (CAF19) separately (Figure 6). Notably, GFP-labeled CAFs retained viability at up to 0.2 µM for simvastatin-Cy5.5 conjugates and 2 µM for pravastatin-Cy5.5 conjugates (Figure 6a), while these concentrations inhibited the growth of Panc1 cells. Post-treatment viability results confirmed that statin-dye conjugates selectively killed Panc1 cancer cells at 0.2 μM for simvastatin-Cy5.5 and 2 and 5 μM for pravastatin-Cy5.5 (Figure 6b), demonstrating cancer cell-targeted cytotoxicity with minimal impact on stromal cells. This suggests a promising and novel approach for treating PDAC tumors, which frequently harbor *KRAS* mutations.

**Figure 6.**
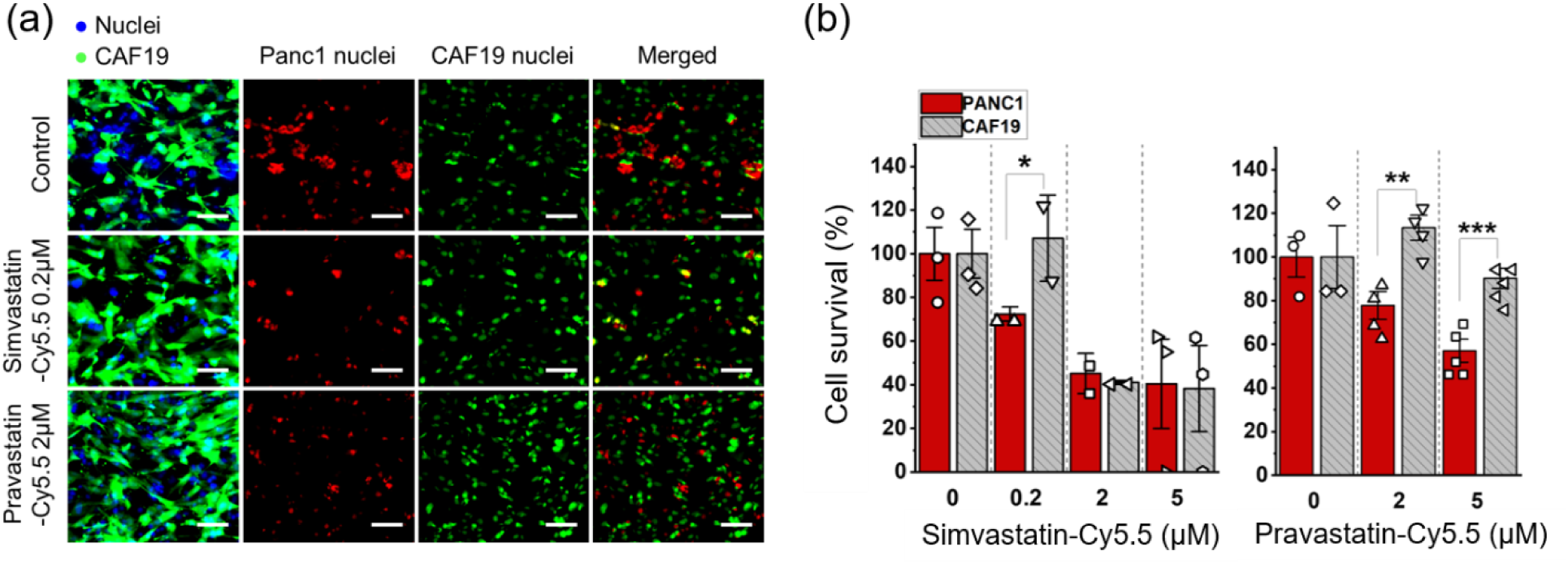
The tumor-selective cytotoxicity of statin-dye conjugates in *in vitro* T-MOC platforms. (a) Fluorescence micrograph showing the co-cultured Panc1 (KRAS^MUT^) and green fluorescent-labeled CAF19 (KRAS^WT^) cell nuclei in the T-MOC device, both with and without treatment of simvastatin-Cy5.5 (0.2 µM) and pravastatin-Cy5.5 (2 µM). CAF19 cells are represented in green, while the nuclear areas of both Panc1 and CAF19 cells appear in blue. The overlapping regions of the nuclear area with the green fluorescent signal are attributed to CAF19 nuclei (cyan), while the remaining nuclear areas correspond to Panc1 nuclei (red). The scale bars represent 100 μm. (b) Cell survival (%) measured after treating the co-cultured Panc1 and CAF19 cells in the T-MOC with simvastatin-Cy5.5 and pravastatin-Cy5.5. Bars represent mean ± S.E. (n ≥ 3). Statistically significant differences are indicated by * for p < 0.05, ** for p < 0.01, and *** for p < 0.001 (student t-test).

## DISCUSSION

In this study, we synthesized statin-dye conjugates by attaching the fluorescent dye Cy5.5 to two commercially available statins, simvastatin and pravastatin, to evaluate their potential for selectively targeting *KRAS*^MUT^ cancer cells. Statins, commonly prescribed for cholesterol reduction, have also shown selective cytotoxicity against *KRAS*^MUT^ cells, leading us to explore whether they might also exhibit selective uptake in these cells. With Cy5.5 labeling, we aimed to track the internalization of statin-dye conjugated specifically in *KRAS*^MUT^ cells and examine their uptake mechanism. To our knowledge, this study represents the first direct examination of statin uptake in *KRAS*^MUT^ cells using fluorescent labeling, offering new insights on statin-based drug delivery into selective targeting potential.

The conjugates demonstrated preferential uptake by *KRAS*^MUT^ cells over *KRAS*^WT^ cells, including various isogenic cell pairs with *KRAS* G12D and G13D mutations. Our findings indicate that macropinocytosis significantly contributes to the uptake of statin-dye conjugates in *KRAS*^MUT^ cells. Although other mechanisms may be involved, our findings confirm that macropinocytosis plays a major role in this selective uptake of statin-dye conjugates in *KRAS*^MUT^ cells. The use of EIPA, a macropinocytosis inhibitor, substantially reduced the internalization of both statin-Cy5.5 in *KRAS*^MUT^ cells, strongly suggesting that macropinocytosis is a key mechanism facilitating the enhanced uptake of these conjugates in *KRAS*^MUT^ cells.^22^ Additionally, we demonstrated that *PTEN* deficiency, known to elevate macropinocytosis through activation of the PI3K/RAC pathway, further increased the uptake of statin-Cy5.5 conjugates. In MCF10A cells with *PTEN*^KO^, the enhanced uptake of these conjugates was significantly reduced upon treatment with either EIPA or a pan-PI3K inhibitor (Figure S4). Taken together, our data support the potential of exploiting the macropinocytosis pathway in targeting tumors with certain specific genetic lesions.

While our data clearly demonstrate enhanced uptake of statin-dye conjugates in *KRAS*^MUT^ cells (Figure 2), the observed cytotoxicity differences between *KRAS*^MUT^ and *KRAS*^WT^ cells were relatively modest. This discrepancy can be attributed to several factors. Cellular uptake does not always directly translate to cytotoxicity, as the intracellular fate of the internalized compounds such as metabolism, degradation, and efflux can significantly impact their biological activity and overall effect on cell viability. Although *KRAS*^MUT^ cells exhibit increased macropinocytotic uptake, their dependency on the mevalonate pathway may differ from that of *KRAS*^WT^ cells. As a result, even with differential uptake, the downstream cytotoxic effects may be relatively modest between the two cell types.

We initially hypothesized that our statin-dye conjugates might be internalized *via* albumin binding, as statins are known to bind albumin strongly, and albumin is often taken up through macropinocytosis. However, our findings indicated that the uptake of simvastatin-Cy5.5 was albumin-independent, occurring even in the absence of albumin (Figure S5). Furthermore, the presence of physiological levels of bovine serum albumin did not significantly increase the uptake. These observations suggest an alternative mechanism, potentially involving nanoparticle formation. Given the physicochemical properties of statin-dye conjugates, they may self-assemble into nanoparticles, facilitating their macropinocytotic uptake into KRAS^MUT^ cells. Further studies are warranted to characterize the nanoparticle formation of these conjugates under physiological conditions and its impact on their selective accumulation in KRAS^MUT^ cells.

The T-MOC model, which closely mimics the complex TME of PDAC, provided a realistic platform to evaluate the efficacy and selectivity of these conjugates. Within this model, *KRAS*^MUT^ PCCs co-cultured with CAFs exhibited selective uptake of statin-Cy5.5 conjugates, with simvastatin-Cy5.5 and pravastatin-Cy5.5 accumulating 2.5 and 5 times more, respectively, in *KRAS*^MUT^ Panc1 cells than in CAFs. This differential accumulation underscores their potential for precise targeting within the complex environment of PDAC. Notably, in the same T-MOC model, doxorubicin did not exhibit selective uptake in KRAS^MUT^ Panc1 cells compared to CAF19 stromal cells, whereas statin-dye conjugates preferentially accumulated in KRAS^MUT^ cells. This contrast suggests that the physicochemical properties of statin-dye conjugates, potentially including nanoparticle formation and macropinocytosis-driven uptake, differentiate them from conventional small-molecule chemotherapeutics. While doxorubicin relies on passive diffusion and transporter-mediated uptake, statin-dye conjugates may be actively internalized through macropinocytosis, enabling selective tumor targeting. Furthermore, the higher drug uptake affinity (k_on_) observed in Panc1 cells further supports the idea that *KRAS*^MUT^ cells may have enhanced endocytic activity, particularly macropinocytosis. These findings highlight the potential advantages of leveraging macropinocytosis in designing targeted drug delivery strategies for KRAS^MUT^ cancers.

## CONCLUSIONS

Our study shows that synthesizing statin-Cy5.5 conjugates, which are selectively internalized into *KRAS*-transformed tumor cells *via* macropinocytosis, can significantly enhance the selectivity of drug delivery. This finding demonstrates strong potential for targeting *KRAS*^MUT^ and *PTEN*-deficient tumors, where macropinocytosis is upregulated. By leveraging this pathway, statin-based drug conjugates could broaden the application range of existing drugs, creating a novel class of potential treatments for *KRAS*-driven tumors. Future research should focus on optimizing these statin-drug conjugates to achieve selective and effective treatments, which will be crucial for maximizing their potential and possibly extending their use to other cancers with similar characteristics.

## EXPERIMENTAL PROCEDURES

### Synthesis of statin-dye conjugates

Simvastatin-Cy5.5: Cy5.5 acid (15 mg, 0.026 mmol, Lumiprobe, MD, USA) was added to a mixture of simvastatin (11 mg, 0.026 mmol), EDC (5.5 mg, 0.035 mmol), and DMAP (1.6 mg, 0.012 mmol) in dichloromethane (DCM, 1 mL). The mixture was stirred at room temperature for 1 h, followed by solvent evaporation. The residue was purified by silica gel column chromatography using DCM/methanol (19/1) as the mobile phase. The final product was obtained as a blue solid. The mass and purity of simvastatin-Cy5.5 were confirmed by liquid chromatography-mass spectrometry (1260 Infinity II; Agilent Technologies, CA, USA). The calculated m/z was 983.6, with a measured m/z of 983.7. The purity of simvastatin-Cy5.5 was analyzed by HPLC in solvent gradient conditions of acetonitrile/H_2_O from 5:95 to 100:0 for 30 min, followed by 100:0 for 5 min.

Pravastatin-Cy5.5: A mixture of pravastatin (20 mg, 0.045 mmol), EDC (14 mg, 0.090 mmol), and NHS (8.6 mg, 0.075 mmol) in dimethylformamide (DMF) was stirred at RT for 30 min. Subsequently, Cy 5.5 amine (34 mg, 0.045 mmol, Lumiprobe, MD, USA) was added in one portion to the mixture. The mixture was stirred at RT for 2 h, followed by solvent evaporation. The residue was purified by silica gel column chromatography using DCM/methanol (4/1) as the mobile phase. The final product was obtained as a blue solid. The mass of pravastatin-Cy5.5 was confirmed by matrix-assisted laser desorption ionization-time of flight (MALDI-TOF, Voyager DE-STR; Applied Biosystems, CA, USA), with a calculated m/z of 1087.7, and a found m/z of 1087.8. The purity was confirmed by HPLC analysis in solvent gradient conditions of acetonitrile/H_2_O from 5:95 to 100:0 for 30 min, followed by 100:0 for 5 min).

### Cell culture and cell line information

Panc1 (CRL-1469) and BxPC3 (CRL-1687) cell lines were purchased from American Type Culture Collection. Panc1 cell line was cultured in Dulbecco’s Modified Eagle’s Medium (DMEM; Invitrogen, MA, USA) supplemented with 10% fetal bovine serum (FBS; Invitrogen, MA, USA) and 1% antibiotic-antimycotic solution (Thermo Fisher, MA, USA). BxPC3 cells were maintained in Roswell Park Memorial Institute 1640 (RPMI 1640; GenDEPOT, TX, USA) medium containing 10% FBS and 1% antibiotic-antimycotic solution.

The *KRAS*-inducible human pancreatic ductal epithelial cells (HPDE *iKRAS*) cell line was kindly provided by Dr. Allen-Petersen from Purdue University. HPDE *iKRAS* cells were genetically modified to allow for the *KRAS* G12D mutation to be induced by the presence of doxycycline. Details about the modification and characterization of the cell line are described in Tsang et al.^23^ HPDE *iKRAS* cells were maintained in keratinocyte-serum free medium (Invitrogen, MA, USA) supplemented by bovine pituitary extract (0.05 mg/ml), recombinant human epidermal growth factor (5 ng/ml) and L-glutamine. For *KRAS* induction, HPDE *iKRAS* cells were treated with 25 ng/ml doxycycline for 48 h (HPDE *KRAS*^MUT^), whereas the HPDE *iKRAS* cells were treated with a corresponding dimethyl sulfoxide (DMSO) control media for HPDE *KRAS*^wt^ in the normal culture conditions. The *KRAS* induction was processed after the cells were harvested and seeded in the experimental platforms. CAF19 transduced with enhanced GFP was kindly provided by Dr. Melissa Fishel^24^ where the CAF19 cells were originally obtained from Dr. Anirban Maitra at Johns Hopkins University.^25^ CAF19 cells were maintained in DMEM supplemented with 10% (v/v) FBS, 1% (v/v) GlutaMAX™ supplement (Invitrogen, MA, USA), and 100 µg/ml penicillin/streptomycin. *KRAS* isogenic HCT116 and DLD1 cell lines (*KRAS*^WT^ and *KRAS*^WT/G13D^) were kindly provided by Dr. Bert Vogelstein at Johns Hopkins Kimmel Cancer Center to Dr. Jean Zhao’s Lab (Table S1).^26^

The cells were regularly harvested by 0.05% trypsin and 0.53 mM EDTA (Life Technologies, CA, USA) when grown to ∼80% confluency in 25 or 75 cm^2^ T-flasks and incubated at 37°C with 5% CO_2_. Harvested cells were used for experiments or sub-cultured while maintaining them within the 15^th^ passage.

### Cellular uptake assay

The cells were seeded on poly-D-lysine coated coverslips (Neuvitro, WA, USA) at a density of 60-70%. For the inhibitor treatment experiment, cells were pre-incubated with DMSO (as mock) or different signaling inhibitors for 1.5 h and treated with 50 nM statin-dye conjugates for 1 h. The signaling inhibitor used for pre-treatment is EIPA (50 μM, 1.5 h) for macropinocytosis inhibition. After treatments, the cells were repeatedly washed with PBS and fixed with 4% paraformaldehyde. The nuclei were stained with Hoechst 33342 (Thermo Fisher, MA, USA), and the coverslips were mounted on a glass slide with a mounting medium (90% glycerol/ 0.2% n-propyl gallate/ 20 mM Tris, pH 8.0). The fluorescence images were obtained by spinning disk confocal on an inverted fluorescence microscope.

### Cytotoxicity assay

The cytotoxicity of simvastatin-Cy5.5, pravastatin-Cy5.5, and Cy5.5 was determined by 3-(4,5-Dimethylthiazol-2-yl)-5-(3-carboxymethoxyphenyl)-2-(4-sulfophenyl)-2H-tetrazolium (MTS) assay which is a colorimetric method to measure cellular proliferation (CellTiter 96® AQueous One Solution, Promega, WI, USA). Briefly, cells were pre-cultured in a 96-well plate for 48 h to achieve ∼60% confluency. The cells were then treated with varying concentrations (0, 0.05, 0.1, 1, 5, 10 μM) of simvastatin-Cy5.5, pravastatin-Cy5.5, or Cy5.5 for 24 h. For the drug solutions, the induction medium containing 25 ng/ml doxycycline was used for the HPDE *KRAS*^MUT^ cells, while a base medium consisting of 1% DMSO was used for the HPDE *KRAS*^WT^ cells. Following this treatment, cell proliferation was assessed using the MTS solution according to the vendor’s instructions. Viability was quantitatively measured by comparing the relative absorbance values to those of the respective control groups (0 μM) for each cell type.

### Cellular uptake assay using T-MOC model

The uptake of the statin-dye conjugates was assessed in PCC-CAF co-cultured T-MOC model to investigate differential response in drug accumulation between PCC and CAF cells. The co-cultured T-MOC model was a microfluidic platform to demonstrate dynamic transport as we described in our prior studies.^18–21^ Briefly, the T-MOC model is composed of capillary, interstitial, and lymphatic channels. The pravastatin-Cy5.5 is accumulated in the cell body through cellular uptake while the compound transports from capillary to interstitial, then lymphatic channels. The drug transport is governed by the perfusion flow condition of the T-MOC which is controlled by differences in hydrostatic pressure of the capillary, interstitial, and lymphatic channels as described in our prior publications.^19, 20^ In the present study, a hydrostatic pressure difference of 20 mm H_2_O was used to mimic the average interstitial flow rates typically observed in TME.^27^ We measured the accumulation by using a live-cell imaging technique with time-lapse microscopy for fluorescent statin-dye conjugates. An inverted microscope (Olympus IX71, Japan) was equipped with a stagetop incubator as described in,^28^ which allowed maintaining the microfluidic platform at 37°C with 5% CO_2_ environment during imaging. After the 2 μM of each statin-dye conjugate (simvastatin-Cy5.5 and pravastatin-Cy5.5) and free Cy5.5 was introduced into the capillary channel, Cy5.5 fluorescence in the T-MOC platform was captured every 2 h for 24 h-duration. Temporal drug accumulation in the cells was measured by fluorescence intensity at each cell type where the intensity was calibrated with fluorescence of 2 μM of the corresponding compound. The accumulation was measured for each cell type separately. Since the CAF19 cell line is transfected with GFP, CAF19 cell area was defined with the FITC-fluorescence. On the other hand, Panc1 cell boundary was defined by using the bright-field images. The contrast differences between cells and background were processed to convert the images to monochrome by using ImageJ. The control experiment with doxorubicin also followed the above procedure to compare the differential drug accumulation. All experiments were repeated at least 3 times for each treatment group. The data was reported in the form of mean ± standard deviation.

To quantitatively address the cell-type specific accumulation of the statin-dye conjugates, we modeled the drug intracellular transport model based on mass conservation (Figure 5d)^21^.

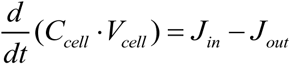

where C_cell_ is an intracellular drug concentration and V_cell_ is an estimated volume of each cellular compartment. Transient drug accumulation is governed by a balance between drug influx (J_in_) and efflux (J_out_) at the cell surface. The cell’s drug uptake affinity (k_on_) at the cell surface and drug concentration at the extracellular region near the cells (C_ex_) determine the drug influx, while the drug efflux is dependent on the drug dissociation affinity (k_off_) and intracellular drug concentration (C_cell_). Assuming that k_on_ and k_off_ are constant parameters, we quantitatively estimated the cellular capability of drug uptake and efflux.

### Cytotoxicity assay using T-MOC model

The efficacy of statin-dye conjugates was assessed in the T-MOC to investigate the differential cell response between PCC and CAF cells. Cells cultured in the T-MOC were pre-cultured for 2 days after loading. Then, simvastatin-Cy5.5 of 0 (control), 0.2, 2, and 5 μM and pravastatin-Cy5.5 of 0, 2, 5 μM was perfused through for 24 h. After the drug treatment, cells were washed with the normal culture medium and cultured in the normal condition for an additional 2 days to have sufficient time to capture the latent effects of drug action. At the final stage of the experiment, cells’ nucleic acid was stained with Hoechst 33342. Although nucleic acid staining marks both live and dead cells, viable cells were assessed in the T-MOC platform on the observation that nuclei of dead cells diminish after 48 hours of post-culture, as verified using Propidium Iodide, dead cell staining in our precious work (Moon, Ozcelikkale et al., 2020). Cell survival was defined by normalizing nuclear area of the treatment group to control group (0 μM) as follow:

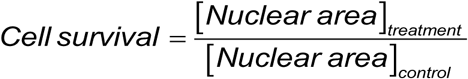

The cell survival metric indicated drug response to the drug by comparing viable cell growth of the treatment group with respect to the growth of control. In the co-culture model, PCCs (Panc1) and CAFs (CAF19) were distinguished by using green-fluorescent CAF19. Specifically, the stained nuclear area included Panc1 and CAF19, while CAF19 nuclear area was identified based on overlap with the green fluorescent signal. While the stained nuclear area includes both Panc1 and CAF19, the CAF19 nuclear area overlaps with the green fluorescent signal. By selectively measuring the nuclear area that overlaps with green fluorescence, we quantified CAF19 nuclei, while the remaining nuclear area was attributed to Panc1 cells (Figure 6a). All experiments were repeated at least 3 times for each treatment group. The data was reported in the form of mean ± standard estimated error (S.E.). Data points in the drug cytotoxicity assay with T-MOC were statistically analyzed by using student t-test. The comparison was done with cell survivals between PCC and CAFs. The differences were recognized as statistically significant when p-value < 0.05.

### Statistical analysis

To compare results, groups were evaluated using the Student’s t-test, with each group containing at least three biological replicates (n ≥ 3). Statistical significance was indicated by p-values, with a threshold of p < 0.05 for significance. We presented the statistical significance of p < 0.05, 0.01, 0.001 and 0.0001 as *, **, *** and **** respectively.

## ASSOCIATED CONTENT

### Supporting Information

The Supporting Information is available.

## AUTHOR INFORMATION

### Corresponding Authors

**Thomas M. Roberts** – *Department of Cancer Biology, Dana-Farber Cancer Institute, Harvard Medical School, Boston, MA, USA*;

**Bumsoo Han** – *School of Mechanical Engineering, Purdue University, West Lafayette, IN, USA; Purdue Institute for Cancer Research, Purdue University, West Lafayette, IN, USA; Department of Mechanical Science and Engineering, Materials Research Laboratory and Cancer Center at Illinois, University of Illinois Urbana-Champaign, Urbana, IL, USA; Chan Zuckerberg Biohub Chicago, Chicago, IL, USA*;

**Ju Hee Ryu** – *Medicinal Materials Research Center, Biomedical Research Institute, Korea Institute of Science and Technology (KIST), Seoul, Republic of Korea; KU-KIST Graduate School of Converging Science and Technology, KIST school, University of Science and Technology, Seoul, Republic of Korea*;

### Authors

**Hye-ran Moon** – School of Mechanical Engineering, Purdue University, West Lafayette, IN, USA; Stem Cell Convergence Research Center, Korea Research Institute of Bioscience and Biotechnology (KRIBB), Daejeon, Republic of Korea

**Zhenying Cai** – Department of Cancer Biology, Dana-Farber Cancer Institute, Harvard Medical School, Boston, MA, USA

**Bo Kyung Cho** – Medicinal Materials Research Center, Biomedical Research Institute, Korea Institute of Science and Technology (KIST), Seoul, Republic of Korea

**Hyeyoun Chang** – Department of Cancer Biology, Dana-Farber Cancer Institute, Harvard Medical School, Boston, MA, USA; Medicinal Materials Research Center, Biomedical Research Institute, Korea Institute of Science and Technology (KIST), Seoul, Republic of Korea

**Seung Taek Hong** – Medicinal Materials Research Center, Biomedical Research Institute, Korea Institute of Science and Technology (KIST), Seoul, Republic of Korea; Division of Biohealthcare, Department of Echo-Applied Chemistry, Daejin University, Gyeonggi-do, Republic of Korea

**Jean J. Zhao** – Department of Cancer Biology, Dana-Farber Cancer Institute, Harvard Medical School, Boston, MA, USA

**Ick Chan Kwon –** Department of Cancer Biology, Dana-Farber Cancer Institute, Harvard Medical School, Boston, MA, USA; Medicinal Materials Research Center, Biomedical Research Institute, Korea Institute of Science and Technology (KIST), Seoul, Republic of Korea; KU-KIST Graduate School of Converging Science and Technology, KIST school, University of Science and Technology, Seoul, Republic of Korea

### Author Contributions

^Δ^H.M., Z.C. and B.K.C. contributed equally to the work.

*T.M.R., B.H. and J.H.R. contributed equally to the work.

### Notes

The authors declare no competing financial interest.

## ACKNOWLEDGMENT

We thank Dr. In-San Kim and Dr. Gi-Hoon Nam for their helpful comments and input. This work was supported by the Intramural Research Program of KIST and the National Research Foundation of Korea (NRF) grant funded by the Korea government (MSIT; No. RS-2024-00463774 to JHR). This work was partially supported by grants from the National Institutes of Health (U01 HL143403, R01 CA254110, U01 CA274304 to BH). HM was partially supported by the Purdue University Center for Cancer Research (P30 CA023168) and a Postdoc Challenge Award from Indiana Clinical and Translational Sciences Institute which is funded in part by Award Number UM1TR004402 from the National Institutes of Health.

## ABBREVIATIONS

PDAC: pancreatic ductal adenocarcinoma
TME: tumor microenvironment
CAFs: cancer-associated fibroblasts (CAFs)
ECM: extracellular matrix
PCC: pancreatic cancer cells
EDC: 1-ethyl-3-(3-dimethylaminopropyl) carbodiimide
DMAP: 4-dimethylaminopyridine
NHS: N-hydroxysuccinimide
DCM: dichloromethane
DMF: dimethylformamide
HPLC: high-performance liquid chromatography
*KRAS*^MUT^: *KRAS* mutation
*KRAS*^G12D^: *KRAS* G12D mutation
*KRAS*^WT^: *KRAS* wild-type
HPDE: human pancreatic epithelial cells
EIPA: 5-(N-ethyl-N-isopropyl)amiloride
*iKRAS*: inducible *KRAS*^G12D^ mutation
*PTEN*^KO^: *PTEN* knockout
*PTEN*^WT^: *PTEN* wild-type
T-MOC: tumor microenvironment-on-a-chip model.

