## Supporting Information for "Statin-dye conjugates for selective targeting of *KRAS* mutant cancer cells"

**SUPPORTING FIGURES**

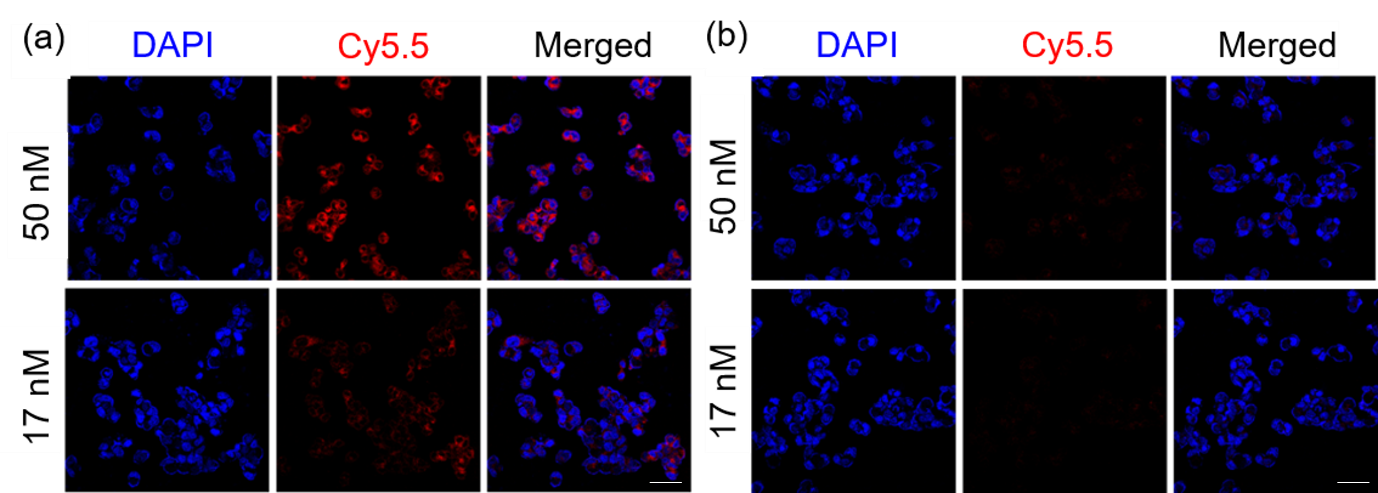

**Figure S1.** Concentration-dependent uptake of simavastatin-Cy5.5 in KRAS-mutant (*KRAS*^MUT^)Panc1 cells. Confocal microscopy images showing the cellular uptake of simvastatin-Cy5.5 (top) and Cy5.5 alone (bottom) in KRASMUT Panc1 cells at two different concentrations: 50 nM and 17 nM. Nuclei was stained with DAPI (blue), and Cy5.5 fluorescence is shown in red. Simvastatin-Cy5.5 exhibited concentration-dependent uptake, with higher fluorescence intensity observed at 50 nM compared to 17 nM. In contrast, Cy5.5 alone showed minimal uptake at both concentrations, suggesting that the statin moiety facilitates selective internalization in *KRAS*^MUT^ cells. The scale bars indicate 500 μm.

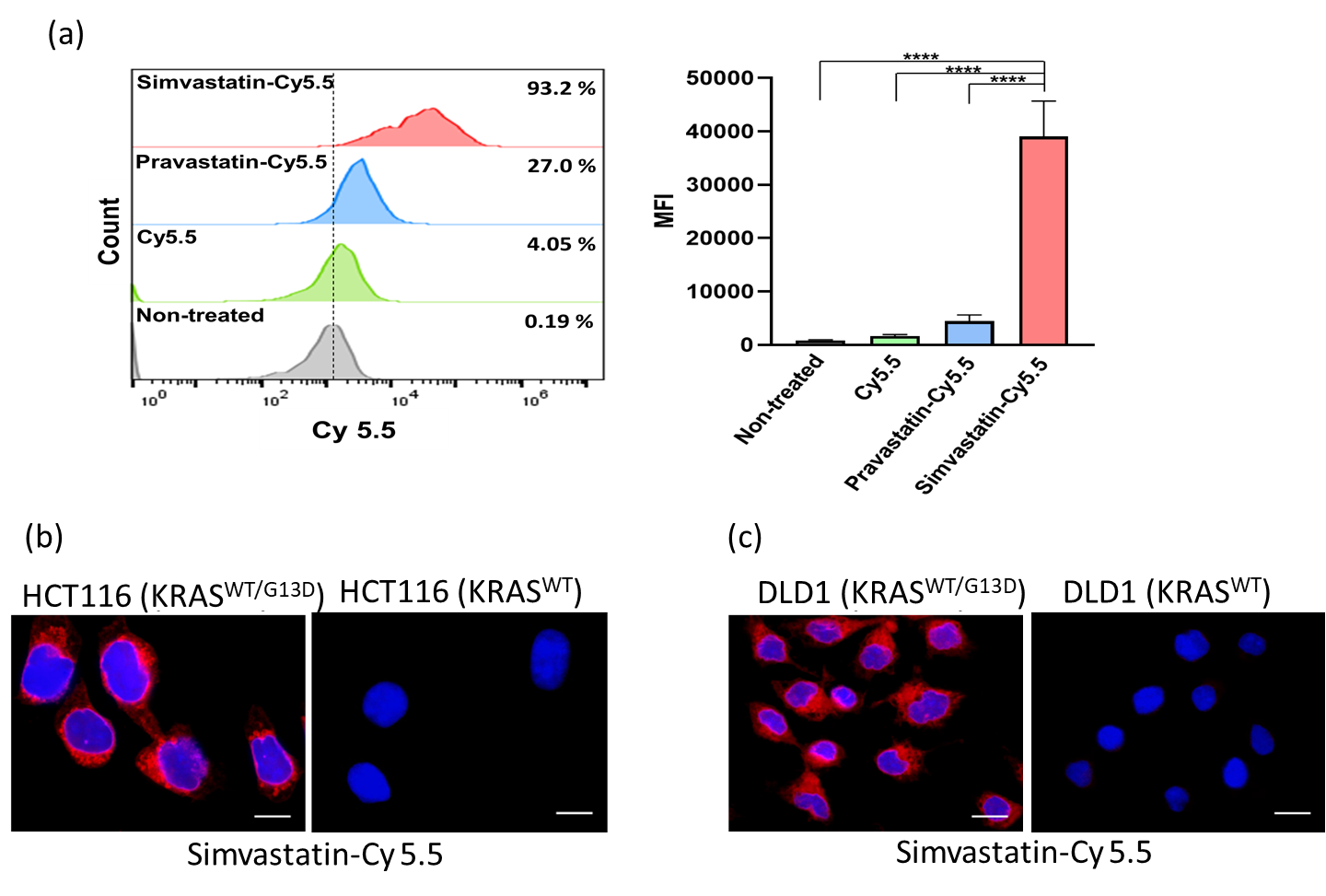

**Figure S2.** (a) Cellular uptake of simvastatin-Cy5.5, pravastatin-Cy5.5, and Cy5.5 in Panc1 cells measured by flow cytometry. Bars indicate Mean ± S.E. (n ≥ 3). Statistically significant differences are represented as **** for p < 0.0001. Cellular uptake of simvastatin-Cy5.5 (red) in isogenic (b) HCT116 and (c) DLD1 cells. The scale bars indicate 100 μm. The cell nuclei were stained with DAPI (blue).

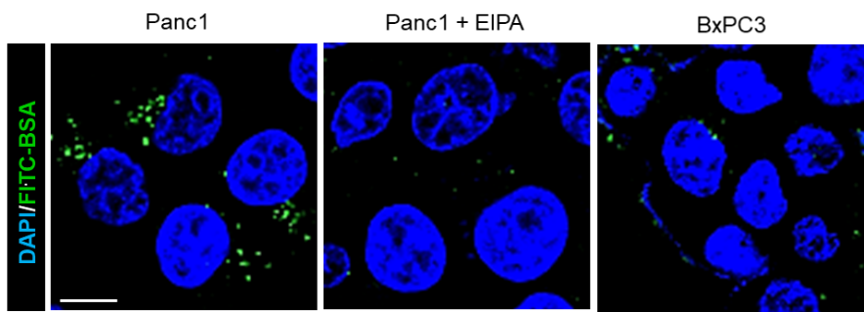

**Figure S3.** Validation of macropinocytosis inhibition by (5-(N-ethyl-N-isopropyl)amiloridein (EIPA; macropinocytosis inhibitor) in *KRAS*-mutant (*KRAS*^MUT^) Panc1 cells. EIPA (50 μM) was pre-treated into cells for 1.5 h, followed by a 1 h incubation with fluorescein isothiocyanate-labeled BSA (FITC-BSA; 2 mg/mL), a widely used marker for macropinocytosis. Representative fluorescence images show FITC-BSA uptake (green) and nuclear staining with DAPI (blue). EIPA treatment significantly reduced FITC-BSA uptake in *KRAS*^MUT^ Panc1 cells, confirming its efficacy in inhibiting macropinocytosis. Additionally, Panc1 cells exhibited higher macropinocytosis activity compared to *KRAS* wild-type (*KRAS*^WT^) BxPC3 cells. The scale bar indicates 100 μm.

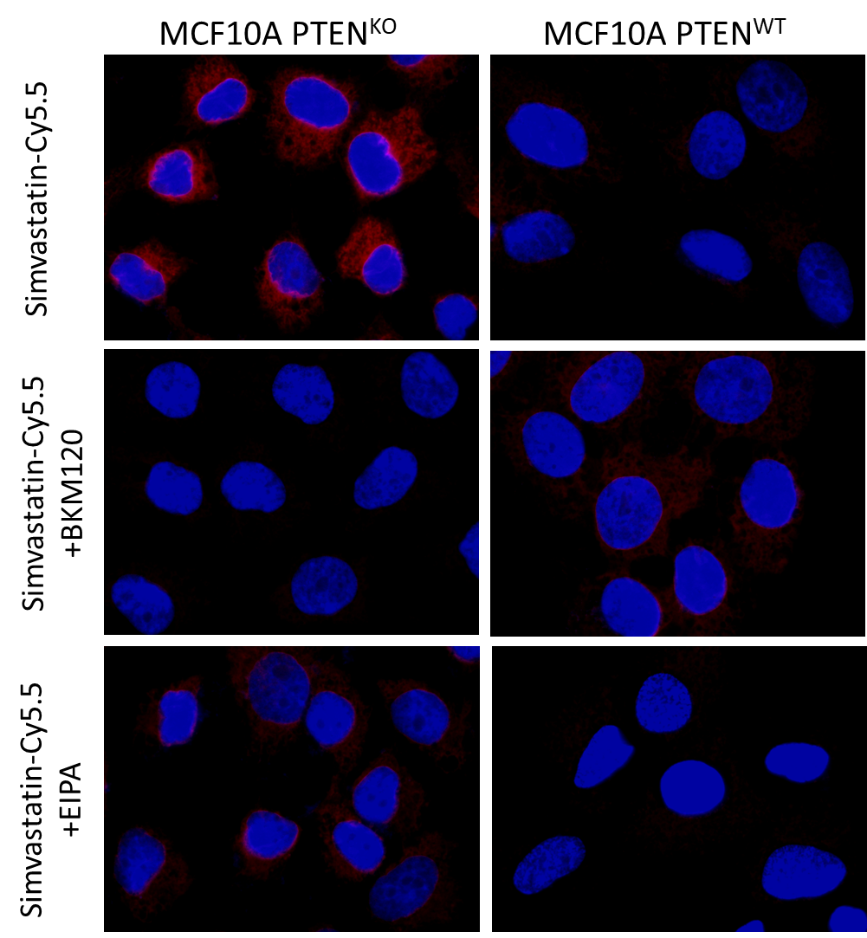

**Figure S4.** Cellular uptake of statin-Cy5.5 conjugates (red) by MCF10A cells with PTEN knockout (*PTEN*^KO^) and wild-type (*PTEN*^WT^). BKM120 (pan-PI3K inhibitor; 10 μM) was pre-treated into cells for 1 h and EIPA (macropinocytosis inhibitor; 50 μM) was pre-treated into cells for 1.5 h. The cell nuclei were stained with DAPI (blue).

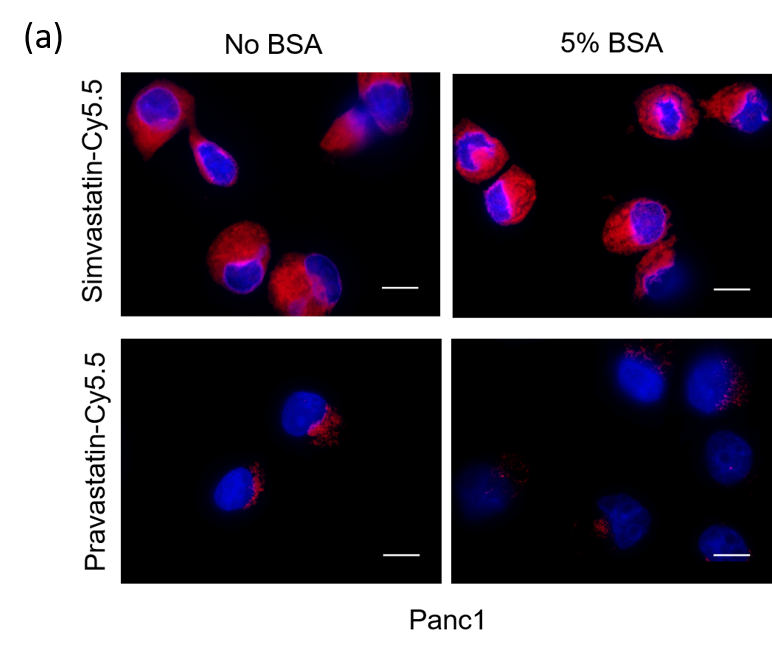

**Figure S5.** Uptake of simvastatin-Cy5.5 (red) and pravastatin-Cy5.5 (red) in Panc1 under no bovine serum albumin (BSA) and 5% BSA condition. The cell nuclei were stained with DAPI (blue). The scale bars indicate 100 μm.

**SUPPORTING TABLE**

**Table S1. Characteristics of the cells used in this study**

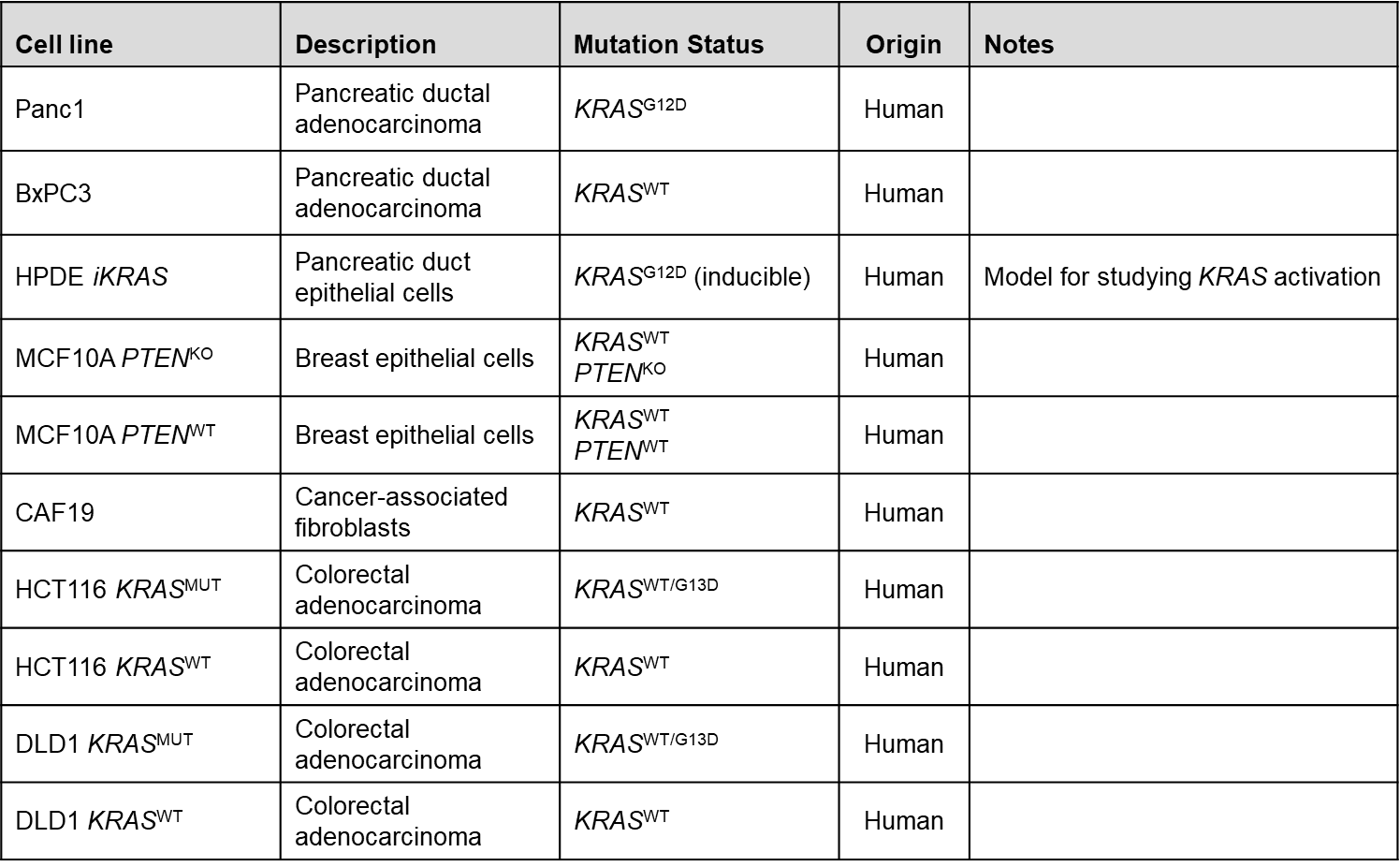
